# Risk of urban Mayaro virus (Alphavirus: Togaviridae) transmission: is *Aedes aegypti* a competent vector?

**DOI:** 10.1101/2023.08.15.553386

**Authors:** Mauricio Daniel Beranek, Octavio Giayetto, Sylvia Fischer, Adrián Luis Diaz

## Abstract

Mayaro virus (MAYV) is an emerging pathogen endemic in Latin America and is the causative agent of fever and polyarthritis. Urban transmission depends on its ability to be transmitted by *Aedes aegypti* and to be amplified by humans. The aim of this study was to evaluate the susceptibility to infection and transmission and the presence of barriers to infection in different populations of *Ae. aegypti* for MAYV. *Ae. aegypti* eggs were collected from Córdoba, Buenos Aires and Rosario Cities (Argentina). Females were infected with five viral loads of MAYV strain (1 to 6 log_10_ PFU/ml) and maintained for 8 days. The presence of infectious viral particles in body, legs, and saliva was detected by plaquing assay in Vero cell monolayers. Through a bibliographic search, *Ae. aegypti* population data from Perú were incorporated and tested with different viral doses of MAYV. We build dose-response curves for *Ae. aegypti* populations to estimate infection (IR), dissemination (DR) and transmission (TR) based on MAYV viral loads detected in humans to estimate transmission risk occurring in an urban environment. The overall IR and DR were significantly associated with the viral doses and were not significantly affected by population origin. We found IR ranging for 3 to 84% (ID50% were higher than 5.5 log_10_ PFU/ml) and a DR reached 78% (DD50% higher than 6.0 log_10_ PFU/ml). The percentage of dissemination based on the infected mosquitoes ranged from 60 to 86% while the percentage of transmission based on disseminated mosquitoes ranged from 11 to 60%. Our results indicate that *Ae. aegypti* populations are not competent vectors for MAYV because they need higher viral doses than those developed by humans (3.9 – 4.5 log_10_ PFU/ml) to become infected. Only a very low proportion of infected mosquitoes with high 5 log_10_ PFU/ml are capable of transmitting it.

## Impacts

- *Aedes aegypti* populations are not competent vectors for MAYV because they need higher viral doses than those developed by humans to become infected and transmit the MAYV.
- Only a very low proportion of infected mosquitoes with high viral doses that developed for humans are capable of transmitting it.

## Introduction

Several arboviruses have emerged and re-emerged causing public health concerns in recent decades (Marcondes et al., 2017). Most arboviruses belong to the families *Bunyaviridae, Flaviviridae*, and *Togaviridae*, and are transmitted mainly by mosquito species of the genera *Aedes* and *Culex* (Gould et al., 2017; Marcondes et al., 2017). *Aedes*-borne arboviruses [yellow fever virus (YFV), dengue virus (DENV), chikungunya virus (CHIKV), and Zika virus (ZIKV)] are worldwide spread and of maximum public health concern because of their potential for urbanization (Gould et al., 2017; Marcondes et al., 2017).

Mayaro virus (MAYV; *Alphavirus*, *Togaviridae*) is an emerging pathogen endemic in South and Central America (Auguste et al., 2015). It was first isolated from febrile workers in Trinidad & Tobago in 1954 (Anderson et al., 1957). Since then, the virus has caused sporadic outbreaks of febrile disease in Latin America and exported cases have been observed in European and North American countries (Acosta-Ampudia et al., 2018). MAYV infection in humans can cause an acute febrile illness characterized by rash and arthralgia (Tesh et al., 1999). Since these symptoms can be caused by other arboviruses like DENV, CHIKV and ZIKV (Zuchi et l., 2014; Gonzalez-Escobar et al., 2021) it is often misdiagnosed (Acosta-Ampudia et al., 2018). Little is known about the MAYV transmission network. It is currently accepted that MAYV is transmitted in a sylvatic network by *Haemagogus janthinomys* with non-human primates as the primary hosts. Other secondary hosts (birds, wild or domestic mammals, reptiles) and different genera of mosquitoes could be implicated (Acosta-Ampudia et al., 2018; Diagne et al., 2020; Caicedo et al., 2021; Celone et al., 2021). In urban areas, domestic mosquitoes are potential vectors of MAYV that could transmit the virus to humans under relevant ecological conditions (Diagne et al., 2020).

MAYV mainly represents a risk for people who live and work in or nearby tropical forests in the Amazon basin (Pereira et al., 2021). However, recent reports alerted about the urbanization risk of MAYV in urban areas of Latin America (Trinidad & Tobago - 2014, French Guiana - 2020, Perú - 2016, Venezuela - 2010, and Brazil - 2014 (Auguste et al., 2015; Alves Esposito & Lopes da Fonseca, 2017; WHO, 2020; Aguilar-Luis et al., 2021; Gonzalez-Escobar et al., 2021). Previous studies with *Ae. aegypti* (Rockefeller strain) showed MAYV low transmission rate for two strains (BeAr 505411 and BeAn 343102) (Brustolin et al., 2018), however, MAYV were transmitted efficiently (strain IQT4235 and strain TRVL 4675) in populations of *Ae. aegypti* from Perú (Long et al., 2011) and Florida (Wiggins et al., 2018). This discrepancy in vector competence could be due to the genetic differences in the *Ae. aegypti* populations or the viral strain evaluated for the experiment (Long et al., 2011; Brustolin et al., 2018; Wiggins et al., 2018; Diop et al., 2019; Alto et al., 2020).

The risk of urbanization for any arbovirus depends on its ability to be transmitted by urban mosquitoes such as *Ae. aegypti* and to be amplified by humans. That is why *Aedes-*borne viruses spill over urban environments and initiate urban outbreaks frequently (Weaver & Reisen, 2010). The aim of this study was to evaluate the susceptibility to infection and transmission of *Ae. aegypti* for MAYV using a viral dose-response approach to identify barriers to infection and dissemination in different populations.

## Materials and Methods

### Virus stock preparation

The BeAr 20290 MAYV (genotype L) strain isolated from *Haemagogus* sp. mosquitoes in Brazil in 1960 was used (Hoch et al., 1981). Viral stocks were prepared by inoculation of 24 hours VERO cells monolayers (African green monkey kidney, *Cercophitecus aethiops*), harvested on 3rd post-inoculation day and centrifuged at 11,000 g for 30 min. Five serial dilutions of MAYV in minimal essential medium supplemented with 10% fetal bovine serum and 1% gentamicin were used.

### Aedes aegypti populations and mosquitoes rearing

*Ae. aegypti* eggs were collected from the cities of Córdoba (Córdoba Province), Buenos Aires (Buenos Aires Province) and Rosario (Santa Fe Province), Argentina (Figure 1). Eggs were stored at room temperature with stable environmental humidity. To synchronize and maximize hatching, mosquito eggs were immersed in 2 liters of distilled water with fish food and vacuumed for 1 hr. Twenty-four hrs after hatching, mosquito larvae were transferred to plastic rearing pans (22 cm in length, 30 cm in width, and 7.5 cm in height) along with distilled water and fed every day. When pupae developed, they were transferred to cups and placed in insect-rearing cardboard cages (22 cm in diameter and 23.5 cm in height). Mosquitoes were reared under controlled environmental conditions of 12 hr light and 12 hr dark photoperiod, 70% relative humidity, and 28 °C. Adults were held in cages to allow them to mate and provided with 10% sucrose solution in cotton wicks. Adult females were deprived of sucrose for 48 hr before the vector competence experiments.

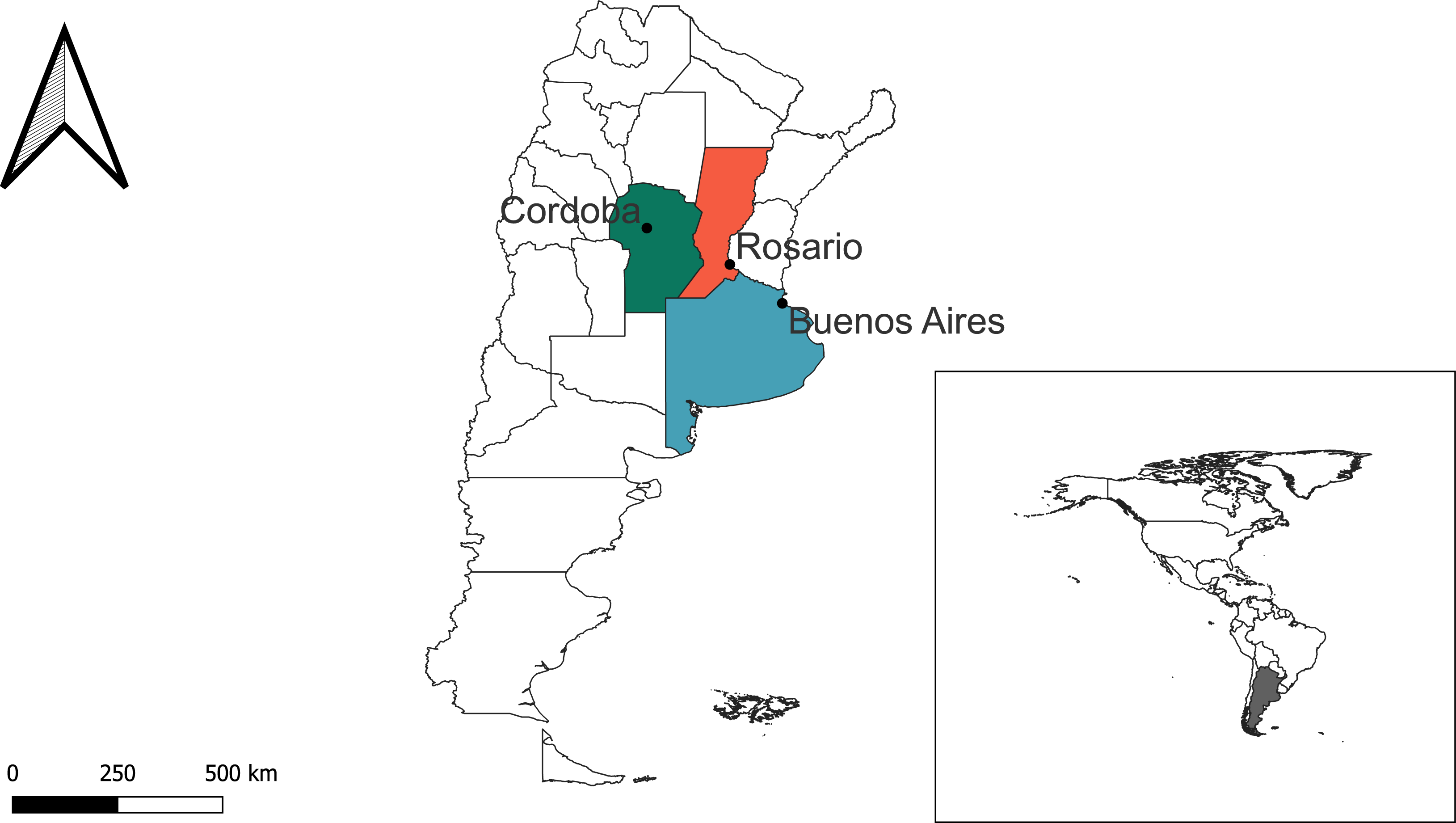

### Mosquito infection

F2 female mosquitoes aged 6–9 days were allowed to feed on MAYV infectious blood meal using an artificial feeding system with hog intestine membranes. Blood meals were prepared as follows: five 10-fold serial dilutions of the 6.5 log_10_ PFU/ml MAYV stock were mixed 1:1 with defibrinated human blood provided by the Blood Bank Foundation (Córdoba, Argentina). 100 µl of each mixture was stored at −80°C for titration. After one hour of exposure, mosquito females were cold anesthetized, and fully engorged mosquitoes were transferred back to their original cages. Not engorged or partially engorged females were discarded. Mosquitoes were maintained under the same conditions previously described.

### Plaquing assay and virus titration

After the incubation period (day 8th after feeding exposure), each virus-exposed mosquito was anesthetized by using triethylamine. Legs and wings were removed, and the proboscis was inserted into a capillary tube with 30 µl glycerin for 30 min for salivary secretion collection. Samples of body and legs were triturated mechanically in a pestle and all samples were centrifuged at 11,000 g for 30 min for clarification. All samples were preserved at –80°C in tubes containing 300 μl phosphate buffer saline (PBS). Salivary secretions were 10-fold serially diluted in PBS for viral titration.

100 μl of each sample were inoculated in a confluent Vero cells monolayer into 24-well plates. The virus was allowed to adsorb for 1 hr at 37°C, an agar overlay was added, and plates were held at 37°C and 5% CO_2_ and observed for cytopathic effect assay over 3 days. After this procedure, 10% formaldehyde (∼0.5 ml) was introduced into each well, agarose plugs were removed, and cells were covered with a crystal violet stain (70% water, 30% methanol, and 0.25% crystal violet) to visualize plaques.

### Data collection and statistical analyses

In order to construct viral dose-response curves for MAYV in other *Ae. aegypti* populations we reviewed original research articles that attempted to identify vector competence in *Ae. aegypti* with different viral doses of MAYV. We first searched PubMed/MEDLINE and Google Scholar for English-language articles. We found 10 scientific papers that referred to the vector competence of *Ae. aegypti* and MAYV but only one paper fulfilled our requirements. This article has information about two *Ae. aegypti* populations collected in Perú (Punchana and Iquitos Districts of Maynas Province) that were tested with different viral doses of MAYV (Long et al., 2011). These data were included in our data analyses and those viral loads were used to estimate the Infection Rate and Dissemination Rate observed in mosquito populations.

We assessed the vector competence of *Ae. aegypti* populations to MAYV by analyzing the infection, dissemination, and transmission rates. We calculated the Infection Rate (IR) as the presence of virus in the body of mosquitoes over the total number of mosquitoes assayed; Dissemination Rate (DR) determined by the presence of virus in the legs of the mosquitoes over the total number of mosquitoes assayed; Transmission Rate (TR) determined by the presence of virus in the saliva of a mosquito over a total number of mosquitoes assayed. Dissemination Effective Rate (DER = proportion of mosquitoes with virus detected in legs among infected mosquitoes); and Transmission Effective Rate (TER = proportion of mosquitoes with virus in the saliva among females with disseminated infection) were calculated in order to identify potential replication barriers in the assayed populations.

By means of binomial error Generalized Linear models, we examined the effects of viral load (continuous variable) and population origin (categorical variable) over the aforementioned calculated rates (IR, DR, TR, DER, TER) (response variables). The significance (α=0.05) of the interactions between viral doses and population origin were assessed using the Likelihood ratio test procedure. To establish the 50% Infectious Dose (ID50) and 50% Dissemination Dose (DD50) for each mosquito population assayed Log-logistic models were used. The 50% Transmission Dose was not estimated since the data obtained was insufficient to perform this analysis. Fitting dose-response curves were also used to estimate infection rate based on MAYV viral loads detected in humans. We searched PubMed/MEDLINE and Google Scholar for English-language articles referring to MAYV loads in humans. We found two scientific papers (Long et al., 2011) and (Pinheiro et al., 1981) that referred to the MAYV human viremic. All statistical analyses were performed in an R 1.4.1717 with the *base* and *drc* packages (Ritz et al., 2015).

### Ethics Statement

The protocol used was approved by the Institute of Virology “*Dr. J. M. Vanella*” (Faculty of Medicine - National University of Córdoba). The human blood was obtained with the BD Vacutainer method. The human blood used in all experiments was obtained from anonymized donors, free of pathogens and collected at the Blood Bank Foundation, and donated to our group for research purposes only.

## Results

Four dose-response curves for MAYV were obtained, three from Argentina and one from Perú. Three *Ae. aegypti* mosquito populations of Argentina were fed with five MAYV concentrations each (ranging from 1.0 to 6.0 log_10_ PFU/ml), and dose-response curves were estimated. This experimental approach allowed us to estimate the proportion of infected (IR) and disseminated (DR) mosquitoes generated at different viral doses. None of the Argentinean mosquito populations fed on infectious doses < 3.5 log_10_ PFU/ml were infected. The minimum IR observed ranged from 5.3% (2/38; 3.9 log_10_ PFU/ml) in the Buenos Aires population, 3.8% (1/26; 3.8 log_10_ PFU/ml) in the Rosario population, 3.1% (1/32; 4.2 log_10_ PFU/ml) in the Córdoba population, and 3% (1/32; 5.57 log_10_ PFU/ml) in the Perú population. The maximum IR observed was 84% (31/37; 7.34 log_10_ PFU/ml) in the Perú population, 65.2% (15/23; 6 log_10_ PFU/ml) in the Rosario population, 41.2% (14/34; 5.6 log_10_ PFU/ml) in the Buenos Aires population, and 22.2% (4/18; 5.7 log_10_ PFU/ml) in the Córdoba population. Regarding the infection dose-response curves, Buenos Aires showed 1.62 more chances of being infected than Perú population, 1.25 than Rosario population and 1.13 than Córdoba population for each logarithm of the viral doses. The overall IR was significantly associated with viral doses (χ^2^ = 48.2 df = 1 p = 3.8 e-12) and was not significantly affected by population origin (χ^2^ = 3.7 df = 3 p = 0.29). The viral dose needed to infect 50% of mosquitoes (ID50) was 5.7 log_10_ PFU/ml (5.24 - 6.23) for Buenos Aires, 5.8 log_10_ PFU/ml (5.44 - 6.19) for Rosario and 6.6 log_10_ PFU/ml for both Córdoba (5.31 - 7.84) and Perú (6.41 - 6.81) (Figure 2). Significant statistical differences were detected in ID50 with the Log-logistic model between Buenos Aires (5.7 [5.24 - 6.23] log_10_ PFU/ml) - Perú (6.6 [6.41 - 6.81] log_10_ PFU/ml) (p = 0.001; t = −3.23) and between Rosario (5.8 [5.44 - 6.19] log_10_ PFU/ml) - Perú (6.6 [6.41 - 6.81] log_10_ PFU/ml) (p = 0.0002; t = 3.71) (Figure 2).

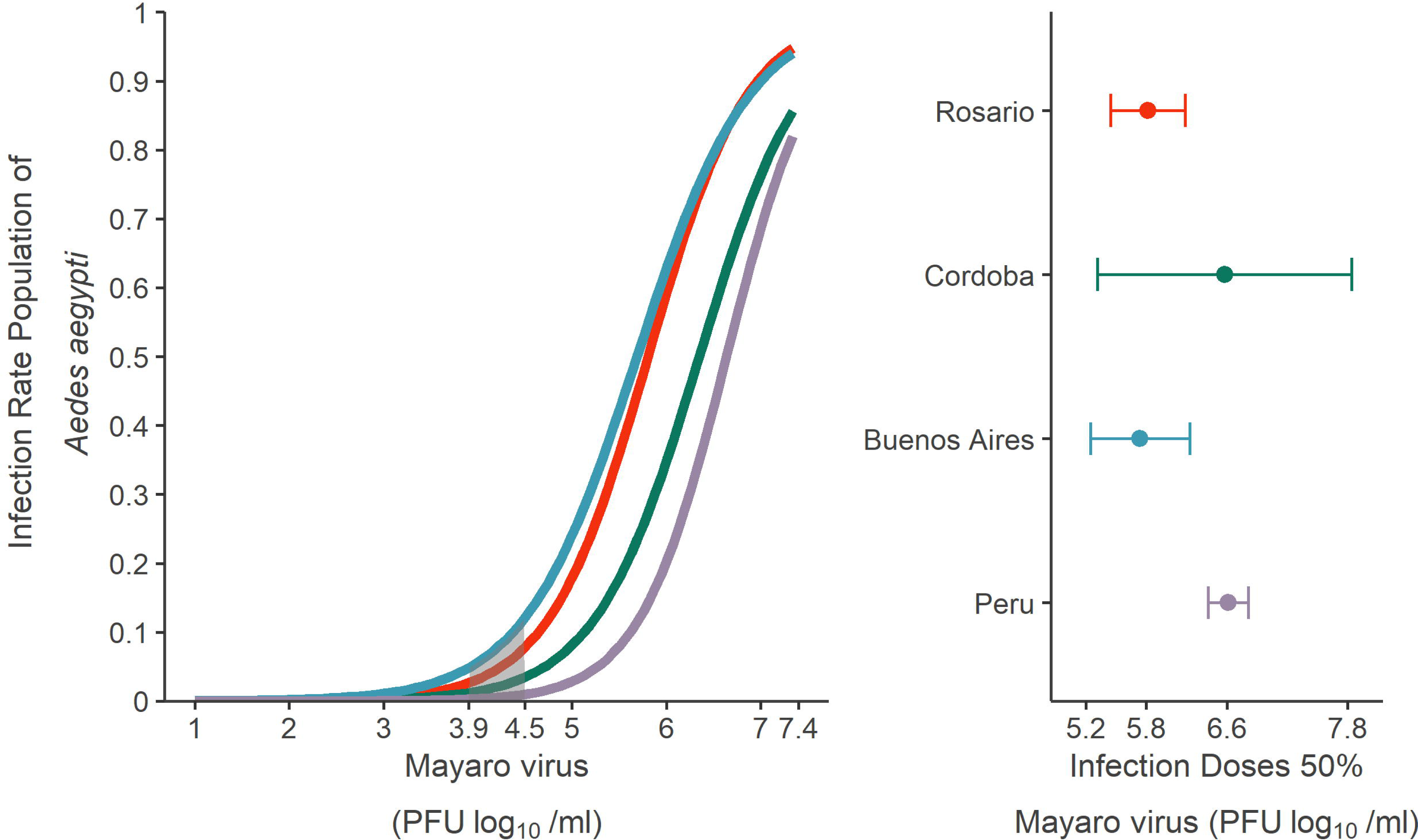

The minimum DR observed ranged from 16.7% (5/30; 5.5 log_10_ PFU/ml) in Rosario population, 8% (2/25; 5.2 log_10_ PFU/ml) in Córdoba population, 3.12% (1/32; 5.57 log_10_ PFU/ml) in Perú population, and 2.6% (1/38; 3.9 log_10_ PFU/ml) in Buenos Aires population. The maximum DR observed was 78.38% (29/37; 7.34 log_10_ PFU/ml) in Perú population, 39.1% (9/23; 6 log_10_ PFU/ml) in Rosario population, 35.3% (12/34; 5.6 log_10_ PFU/ml) in Buenos Aires population and 16.7% (3/18; 5.7 log_10_ PFU/ml) in Córdoba population. Regarding the dissemination dose-response curves, Buenos Aires showed 2.1 more chances of being disseminated than Córdoba population, 1.95 than Rosario population and 1.27 than Perú population for each logarithm of the viral doses. The DR was also positively affected by viral doses (χ^2^ = 29.21 df = 1 p = 6.49e-08) but not by population origin (χ^2^ = 1.46 df = 3 p = 0.69). The viral dose needed to disseminate the viral infection in the 50% of mosquitoes (DD50) was 6.0 log_10_ PFU/ml (5.39 - 6.53) for Buenos Aires, 6.2 log_10_ PFU/ml (5.70 - 6.66) for Rosario, 6.4 log_10_ PFU/ml (5.24 - 7.49) for Córdoba, and 6.7 log_10_ PFU/ml (6.53 - 6.96) for Perú populations (Figure 3). Significant statistical differences were detected in DD50 with the Log-logistic model between Buenos Aires (6.0 [5.39 - 6.23] log_10_ PFU/ml) - Perú (6.7 [6.53 - 6.96] log_10_ PFU/ml) (p = 0.011; t = −2.53) and between Rosario (6.2 [5.70 - 6.66] log_10_ PFU/ml) - Perú (6.7 [6.53 - 6.96] log_10_ PFU/ml) (p = 0.035; t = 2.11) (Figure 3).

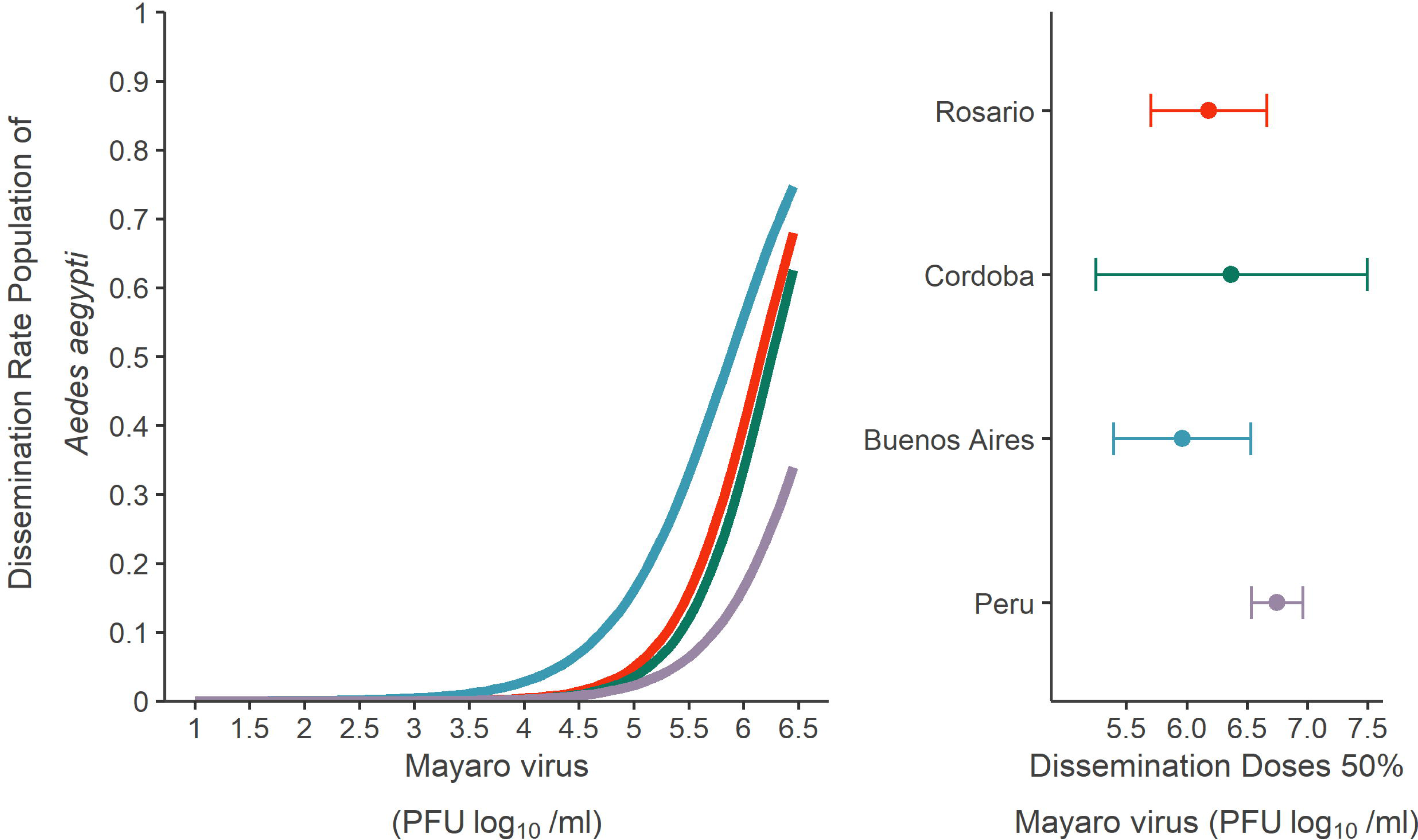

The TR detected in all populations were low and ranged from 4% (1/25) (Córdoba population) to 5.9% (2/34) (Buenos Aires population). Virus titers in saliva varied from 1.3 to 2.2 log_10_ PFU/ml. Infectious viral particles were observed only in the highest viral doses (5.6 log_10_ PFU/ml - Buenos Aires and 6.0 log_10_ PFU/ml - Rosario population) except the Córdoba population in which transmission was observed in the medium viral dose (5.2 log_10_ PFU/ml). Another parameter could not be calculated due to the few saliva samples that were positive for MAYV.

To assess the contribution of *Ae. aegypti* mosquito to initiate an urban MAYV transmission focus, we estimated the transmission threshold in association with viral loads detected in humans. Only 2 scientific articles have information regarding human viremias for MAYV. Long et al. (2011) reported viremias from 22 patients from Perú and Bolivia ranging from 2.7 to 5.3 log_10_ PFU equivalents/ml (PFUe/ml), with a geometric mean of 4.2 log_10_ PFUe/ml (95% confidence interval [CI] = 3.9 – 4.5 log_10_ PFUe/ml). Pinheiro et al. (1981) documented viremia values detected in 21 patients from Brazil. The highest viremia titer was 5 log_10_ PFU/ml (ranging from 1 to 5 log_10_ PFU/ml). Geometric mean and CI cannot be calculated because individual patient data are not available. According to these values and based on our estimated models, human viremias reported are not enough to produce infectious *Ae. aegypti* mosquitoes since their transmission threshold was higher than 5 log_10_ PFU/ml and the minimum infection and dissemination threshold were around 4 log_10_ PFU/ml and close to 5 log_10_ PFU/ml, respectively.

Finally, to measure the presence of barriers along the viral infection process in the *Ae. aegypti* populations, we calculated DER and TER respectively. The percentage of dissemination based on infected mosquitoes ranged from higher than 60% to 85.7%. The highest DER detected were 85.7% (12/14; 5.6 log_10_ PFU/ml) in the Buenos Aires population, followed by Perú population with 75% (6/8; 5.58 log_10_ PFU/ml), Córdoba population with 71.4% (5/7; 5.45 log_10_ PFU/ml), and finally Rosario population with 60% (9/15; 6.0 log_10_ PFU/ml) (Figure 4). The percentage of transmission based on disseminated mosquitoes ranged from 11.1% to 60%. The highest TER detected was in the Perú population [60%; 3/5 (5.58 log_10_ PFU/ml)], and lower in Córdoba [20%; 1/5 (5.45 log_10_ PFU/ml)], Buenos Aires [16.7%; 2/12 (5.6 log_10_ PFU/ml)] and Rosario population [11.1%; 1/9 (6.0 log_10_ PFU/ml)] (Figure 4).

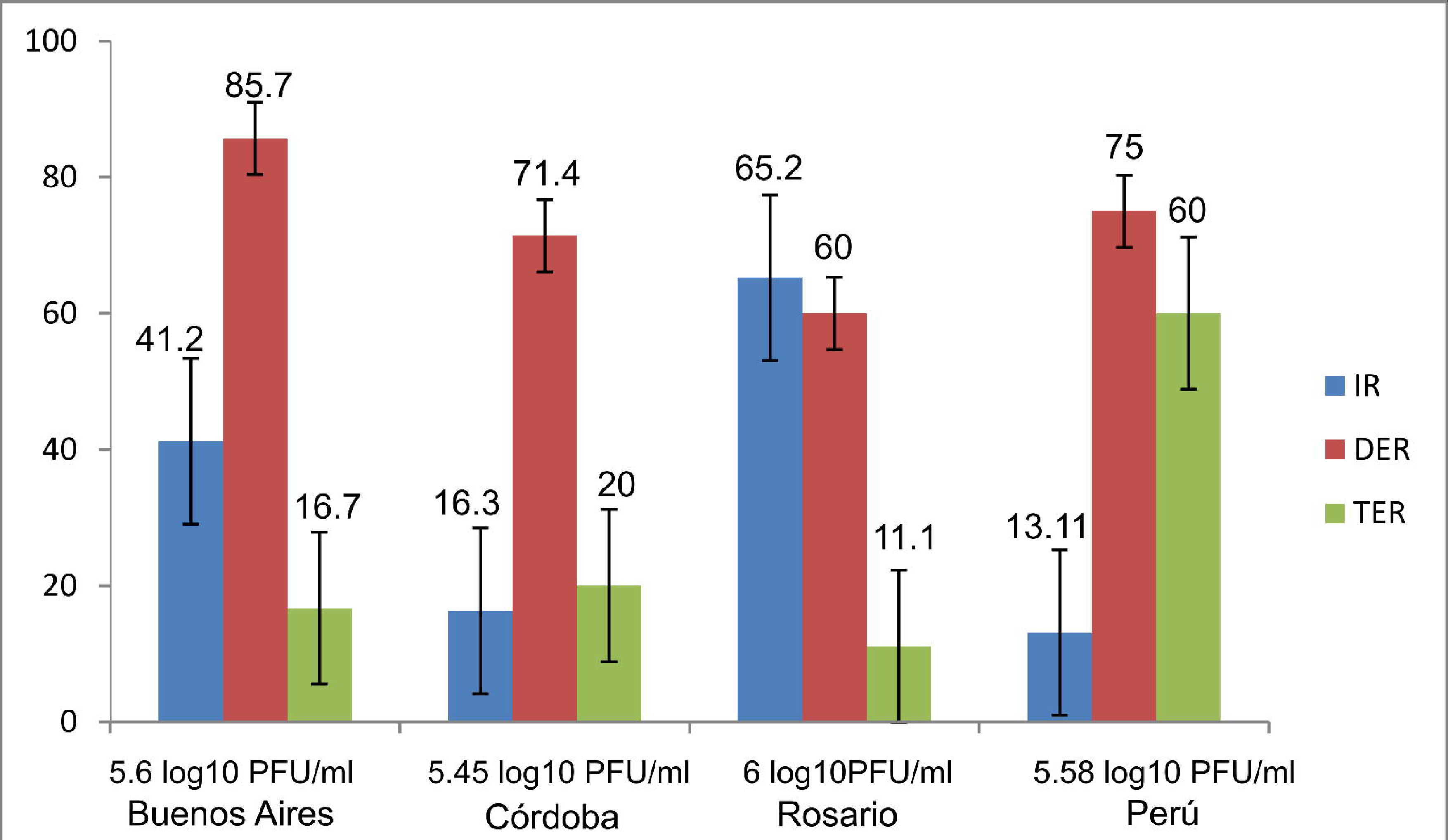

## Discussion

Previous studies have shown that *Ae. aegypti* is a competent vector of MAYV (Long et al., 2011; Brustolin et al., 2018; Wiggins et al., 2018; Diop et al., 2019; Alto et al., 2020; Pereira et al., 2020; Sucupira et al., 2020, Alomar et al., 2022; Krokovsky et al., 2023). Most vector competence studies are performed with limited viral doses and generally use higher viral doses in order to increase the likelihood to detect infected mosquitos. For example, previous studies with higher viral doses of MAYV, showed a high IR of > 90% (9.0 log_10_ PFU/ml) using *Ae. aegypti* populations from Brazil (Sucupira et al., 2020) and IR > 80% of studies using different populations of *Ae. aegypti* (9.0 log_10_ PFU/ml) from Brazil (Pereira et al., 2020), (7.3 log_10_ PFU/ml) from Perú (Long et al., 2011), (7.5 log_10_ PFU/ml) from the USA (Wiggins et al., 2018) and Rockefeller strain with 7.0 log_10_ PFU/ml (Brustolin et al., 2018). These studies offered useful data, but not enough neither to estimate infectious and transmission thresholds nor to evaluate the real ability of *Ae. aegypti* to transmit MAYV. Our study used a viral dose-response curve approach so we were able to estimate infection, dissemination and transmission thresholds in four *Ae. aegypti* populations (three from Argentina and one from Perú). *Ae. aegypti* populations (IR and DR) showed a positive association with the viral doses of MAYV and were not affected by population origin (Figures 2 and 3). Infection, dissemination and transmission of MAYV were detected only at higher viral loads (from 5.2 to 6.0 log_10_ PFU/ml), thus the MAYV infection process was dose-dependent. A dose-dependent effect has been associated with escape barrier from the midgut/saliva gland and occurred only when low doses of the virus had been ingested, thus increasing blood meal viral titers reduced the midgut/saliva gland escape barrier effect (Franz et al., 2015; Kramer & Ciota, 2016; Lim et al., 2018; Merwaiss et al., 2021; Sanchez-Vargas et al., 2021). Similar trends were observed for dissemination rate in *Ae. aegypti* populations. In fact, our results show a high minimum infection threshold (ranging from 3.8 to 4.2 log_10_ PFU/ml). The population origin of *Ae. aegypti* was not a significant factor influencing infection and dissemination of MAYV, however, we observed a higher susceptibility to infection and disseminated infection of MAYV in *Ae. aegypti* from Buenos Aires and Rosario compared to *Ae. aegypti* from Córdoba and Perú reported in the current study. The differences may be associated with virus-specific responses in mosquito and the genetic structure of the mosquito population (Franz et al., 2015; Gloria-Soria et al., 2016). In a previous study, the authors found that the Brazilian population of *Ae. aegypti* is 40% more susceptible to MAYV infection and exhibits 40% higher disseminated infection than the Florida population of *Ae. aegypti* (Alto et al., 2020). Moreover, our study and Long et al. (2011) studies found similar results with different MAYV strains. On the other hand, Brustolin et al. (2018), observed in *Ae. aegypti* that the genotype L exhibited significantly higher infection rates (86.2% at 7 dpi and 51.7% at 14 dpi) compared with the genotype D strain (7.1% at 7 dpi and 0% at 14 dpi).

The susceptibility of a population is generally defined as the concentration of virus required to infect of individuals in the mosquito population (Lounibos et al., 2016). In our study, the ID50% and DD50% were higher than 5.5 log_10_ PFU/ml and 6.0 log_10_ PFU/ml respectively. Unfortunately, we were not able to estimate TD50% since TR detected among populations were low and only the highest viral doses of MAYV produced infectious mosquitoes. However, our data suggested that *Ae. aegypti* mosquitoes cannot transmit the virus if ingested less than 5 log_10_ PFU/ml. The small numbers of positive saliva samples detected could be biased by the microcapillary tube technique used to collect saliva or the presence of a salivary gland infection barrier (SGIB) and salivary gland escape barrier (SGEB) (Gloria-Soria et al., 2016). MAYV titers in saliva varied from 1.3 to 2.2 log_10_ PFU/ml. Similar ranges of saliva titer were obtained for Long et al. (2011) and Wigging et al. (2018). If this is the case, then infectiousness of MAYV in humans should be extremely high in order to produce a productive infection. A previous study found that *Ae. aegypti* salivates an average of 4.7 μg during the blood-feeding process (Devine et al., 1965) but the amount of viral particles transmitted by a vector is highly variable depending on the technique used for detection (Colton et al., 2005; Dubrelle et al., 2009; Anderson et al., 2010). A recent study proposes the presence of virus in mosquito legs could be a more accurate predictor of transmission than the viral detection method using forced salivation into a capillary tube (Gloria-Soria et al., 2022). Even if this were the case, in our study the DR was less than 40% and needed viral doses higher than those developed by human viremias to transmit the virus.

We determined the DER as an estimator of the potential presence of barriers in the midgut of *Ae. aegypti* populations. The studied populations of *Ae. aegypti* showed a DER that ranged from 85.7% to 60%. The populations of *Ae. aegypti* might have an important midgut infection barrier (MIB) that helps prevent viral infection and dissemination. Thus for MAYV, it could be more difficult to infect and enter the cells of the midgut, but once inside, the virus would escape the hemocoel more easily and have a low resistance to the midgut escape barrier (MEB). For example, in female mosquitoes exhibiting an MEB, the virus infected the midgut but was unable to disseminate to other organs, remaining sequestered in the infected midgut cells (Kramer et al., 1981). The presence of MEB has been documented in a variety of mosquito species such as the Lacrosse virus in *Aedes triseriatus* (Grimstad et al., 1985), western equine encephalitis virus in *Culex tarsalis* (Kramer et al., 1981) and Rift Valley fever virus in *Aedes vexans* (Hartman et al., 2021). Once MAYV enters the midgut, infects and replicates in midgut epithelial cells, the virus escapes to the hemocoel from where it disseminates to various organs including salivary glands. We determined the TER as an estimator of the potential presence of barriers in the salivary gland of *Ae. aegypti* populations. The Perú population performed a high TER of 60% meanwhile Argentinean *Ae. aegypti* populations showed a lower TER that ranged from 11.1% to 20%, suggesting the presence of SGIB/SGEB barriers. Further studies are needed on MAYV dynamics of infection/transmission barriers in *Ae. aegypti* populations from Argentina.

Unlike CHIKV, MAYV viremia in humans is short lasting around 5 to 7 days and low (median titters < 5 log_10_ PFU/ml) (Tesh et al., 1999; Long et al., 2011). Previous vector competence studies have demonstrated that vectorial transmission of MAYV requires high levels of viremia (Long et al., 2011; Brustolin et al., 2018; Wiggins et al., 2018; Pereira et al., 2020). In our study, *Ae. aegypti* did not show a high susceptibility to infection nor to transmission (IR from 3% to 84%, DR from 2% to 78% and TR was slightly greater than 4%). Moreover, the minimum transmission threshold was estimated to be higher than 5 log_10_ PFU/ml. This high transmission threshold in *Ae. aegypti* populations could be preventing the development of MAYV human outbreaks in the Americas. However, adaptive mutations in emerging virus strains may affect the intensity of transmission by one species of mosquito and not another (Kramer, 2016) as happened with the emergence of CHIKV and its adaptation to a new mosquito vector *Aedes albopictus* (Tsetsarkin et al., 2007). Recently, Cereghino et al. (2023) observed that T179N mutation in E2 protein increases the vector competence of *Ae. aegypti* to MAYV but decreases pathogenicity and viremia levels in mice model. Mutations at this site could represent potential first steps towards a future urbanization event of MAYV.

Most arboviruses are maintained in enzootic transmission networks and those capable of being vectored by urban mosquitoes such as *Ae. aegypti* and *Ae. albopictus* and being amplified by humans can spill over urban environments and initiate urban outbreaks. This mechanism is one of the most important driving arbovirus emergence worldwide (Weaver & Reisen, 2010; Lwande et al., 2015). According to our data, the tested *Ae. aegypti* populations have low vector competence status for transmission of MAYV as suggested by its relatively low IR and DR, low TR, and high transmission threshold indicating the presence of MIB and SGIB/SGEB barriers. This poor vector competence may be preventing the development of MAYV outbreaks in human settlements in the Americas.

## Acknowledge

Our special thanks to Ambar Canabal Drisun for her support in the field collection of *Ae. aegypti* eggs in Rosario City, Sebastián Blanco for human blood source, Kristin Sloyer for the contribution to MAYV bibliography, and Agustín Quaglia for help in stat analysis.

## Funding information

This research received funding from Ministerio Nacional de Ciencia y Tecnología de Argentina (PICT 2018-1172) and Secretaría de Ciencia y Tecnología–Universidad Nacional de Córdoba (SECyT Consolidar 2018, Universidad Nacional de Córdoba).

## Conflicts of interest

The authors report there are no competing interests to declare.

## Data availability statement

The data that support the findings of this study are available from the corresponding author upon reasonable request.

## Notes

### Competing Interest Statement

The authors have declared no competing interest.

